# orthoCapture: Facilitating Gene Capture Probe Design for Non-Model Species

**DOI:** 10.1101/703942

**Authors:** M. Elise Lauterbur

## Abstract

In non-model species, targeted gene capture (selective enrichment of specific genomic regions of interest) applications in molecular ecology have been limited by the practicalities of capture design. Currently, the minimal requirement for designing capture probes is a transcriptome, or established reference genome for the species of interest. When an established, annotated reference genome is unavailable, one common approach is to design probes from annotated reference genomes (or transcriptomes) of related species. Unfortunately, as divergence between probes and the genome of interest increases, such as occurs during directional selection, capture performance decreases. Here I introduce orthoCapture, a tool to overcome such limitations by mining unannotated whole-genome sequence (WGS) data from non-model species and/or their close relatives to allow probe design using multiple genomic sources. orthoCapture finds orthologs in WGS data from multiple related species to create a set of exon sequences that encompasses the diversity of the exons of interest. These “design sequences” can then be used to design capture probes for the species of interest. orthoCapture thus eliminates the need for transcriptome or whole-genome sequencing for bait capture experiments, making this technique accessible for molecular ecology and conservation studies. Use of orthoCapture is via command-line interface on Unix systems, and requires the input of a gene sequence from an unrelated annotated genome and a fasta database from a target, unannotated genome (e.g., whole-genome shotgun contigs). The output, sequence templates from the nonannotated genomic data, allows probe creation by any commercial company providing gene capture services.

## Introduction

Targeted gene capture (also called targeted enrichment, exon capture, hybrid enrichment, and sequence capture) has become an important method for facilitating the sequencing of targeted sets of genes and conserved elements for evolutionary studies. This method allows the selective enrichment and sequencing of genomic regions of interest by targeting these regions with a set of DNA or RNA probes prior to sequencing. Gene capture balances cost with the number of genes to be sequenced, allowing comparisons among many species or individuals when it is not necessary to acquire whole-genome data (Jones and Good 2016). This is especially relevant since it remains expensive to create whole-genome assemblies for many individual species in a single study, despite recent advances in Next-Generation Sequencing technology and associated cost reductions. Thus, targeted gene capture is especially useful for comparative phylogenetic studies, and it holds potential for broad molecular ecology and conservation applications.

Gene capture applications in molecular ecology and conservation have so far been more limited than in phylogenetics because of the challenge of designing appropriate probes when a high quality, annotated reference genome does not already exist (Jones and Good 2016). Since the probes (also called baits) are intended to hybridize with, and thus “capture,” the regions of interest, they must have nucleotide sequences that complement these regions. When the nucleotide sequence of the regions of interest is known, for example when an annotated reference genome or transcriptome for the species exists, probe design is trivial. However, many species of evolutionary and conservation interest still lack well-annotated reference genomes.

One common approach is to design probes from annotated reference genomes (or transcriptomes) of related species (eg. Cosart et al. 2011; George et al. 2011; Nadeau et al. 2012; Ilves and López-Fernández 2014; Bragg et al. 2016; Jones and Gomulkiewicz 2012; Portik, Smith, and Bi 2016). Unfortunately, as divergence between the probes and the genome of interest increases, capture performance decreases, such that probes perform poorly on regions of interest that are more than 5-12% diverged from the genome used to create the probes (Vallender 2011; Bi et al. 2012; Hancock-Hanser et al. 2013; Hedtke et al. 2013). For example, Vallender (2011) found that capture efficiency in non-human primates dropped off when the genome of interest exceeded 4% sequence divergence from the probes. Hancock-Hanser et al. (2013) found that three-quarters of sequencing gaps (areas with no base calls) occurred in regions of less than 88% sequence identity. It is thus preferable to custom-design probe sets for species of interest (Kadlec et al. 2017).

When well-annotated genomes of the study species are lacking, increasing the diversity of the bait set has been recommended to increase the likelihood of capturing genomic regions of interest (Bi et al. 2012; Schott et al. 2017). This is achieved by designing baits from the reference genomes of multiple, related species. With multiple baits tiled across each region of interest, a bait designed from one species may capture the region even if baits from other species do not (figure 1).

**Figure 1.**
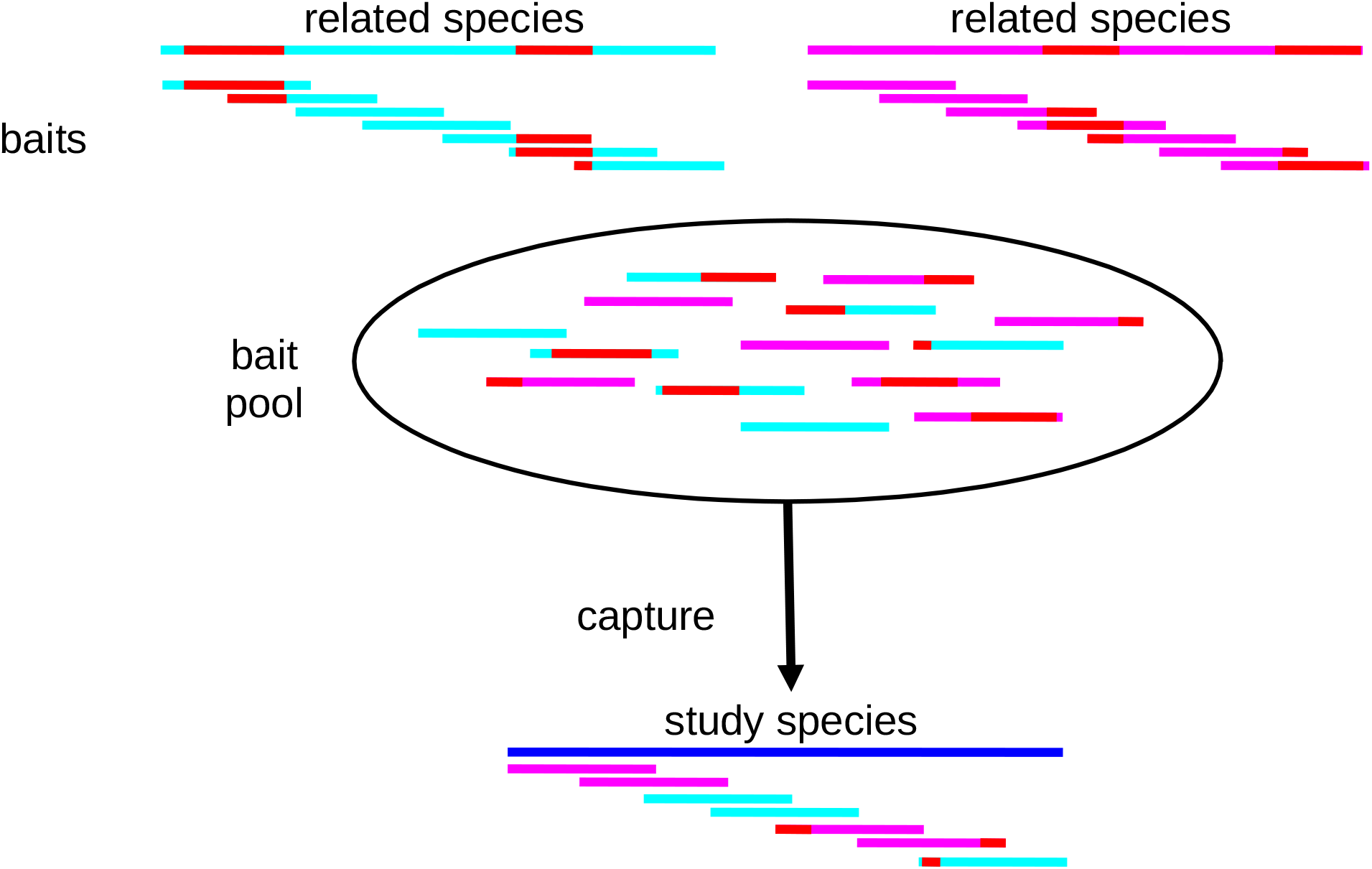
Increased bait (probe) diversity (resulting from baits designed based on multiple related species) increases the likelihood of successful capture of regions of interest from the study species. Dark blue represents the region of interest in the study species. Light blue and purple indicate sequences in the related species orthologous to the region of interest in the study species. Red indicates regions of divergence from the study species. When baits from two or more related species are included in the capture pool, there is increased likelihood of at least one bait matching the sequence in the study species well enough to capture it.

Methods to find and develop bait sets from conserved or ultra-conserved regions for phylogenetic analyses are well-established (Bi et al. 2012; Faircloth et al. 2012; Lemmon, Emme, and Lemmon 2012; Faircloth 2017). These vary in both taxonomic scale and the size of the conserved region to be captured, but the focus remains on producing data for phylogenetic studies rather than studies of molecular evolution. Thus, these methods optimize for easily captured targets instead of coding regions of interest. But in studies of molecular evolution, increased divergence between species is expected, and conserved elements are uninformative about genes under selection. The challenge is to find a balance between baits that are sufficiently similar to the regions of interest, and capturing the sequence diversity that can result from differential selection across species (Schott et al. 2017).

Currently, the minimal requirement for custom-designing bait sets without the reference genome is a transcriptome from the species of interest (Matz 2018; Puritz and Lotterhos 2017). A number of well-tested methods exist to design bait sets from transcriptomes, including Hyb-Seq (Weitemier et al. 2014), MarkerMiner (Chamala et al. 2015), and others (Yang and Smith 2014; Kadlec et al. 2017). Many studies have designed custom methods (eg. Ilves and López-Fernández 2014; Mandel et al. 2014; Hugall et al. 2016; Teasdale et al. 2016; Schmickl et al. 2016; Schott et al. 2017; Johnson et al. 2018) unique to the individual study. Transcriptome-based methods, however, require either an existing transcriptome, or flash-frozen fresh tissue or RNA, and funding to generate a transcriptome. This is frequently infeasible, particularly when the study includes many species without existing transcriptomic resources (thus requiring new transcriptomes for each species), and when the species of interest are elusive or highly endangered, thus sources of RNA are hard to come by (Schott et al. 2017). In these cases, it would be preferable to have a bait design method that used available genomic sequences.

The method required must, first, design baits from existing un-annotated genomic sequences (whole-genome contigs) from the species of interest or, when whole-genome contigs are not available, from multiple related species; second, Use annotations from other species regardless of if they are highly (>10%) diverged from the species of interest; and third, provide a set of sequences that can be curated by standard bait design methods.

Here I introduce orthoCapture, a tool to facilitate gene capture probe design for non-model organisms. It works by mining un-annotated genomic data from non-model species and/or their close relatives. By mining multiple whole-genome shotgun assemblies it can increase the diversity of the resulting probe set, increasing the likelihood of capturing regions of interest. To my knowledge, this is the only such publicly-available tool that can prepare sequences for probe design of non-conserved regions that does not rely on transcriptome data for any of its steps. orthoCapture thus eliminates the need for costly transcriptome creation, or whole-genome sequencing and annotation for bait capture experiments, making this technique more accessible for molecular ecology and conservation studies.

Use of orthoCapture is via command-line interface using Python 3, and requires the input of a gene sequence from an unrelated, annotated genome and a fasta database from a target, unannotated genome (whole-genome shotgun contigs, for example). The output, sequence templates from the nonannotated genomic data, allows probe creation by any commercial company providing gene capture services.

## Methods

### Description

orthoCapture uses six steps to find target sequences for probe creation (figure 2). Here, “target” sequence refers to the putative gene sequence from the whole-genome shotgun (WGS) assembly, whereas “design” sequence is used to refer to the sequence from a well-annotated genome used to mine the WGS assembly. orthoCapture requires a list of genes to be captured identified by gene name (e.g., ND1), as well as design sequences of their individual exons based on annotations.

**Figure 2.**
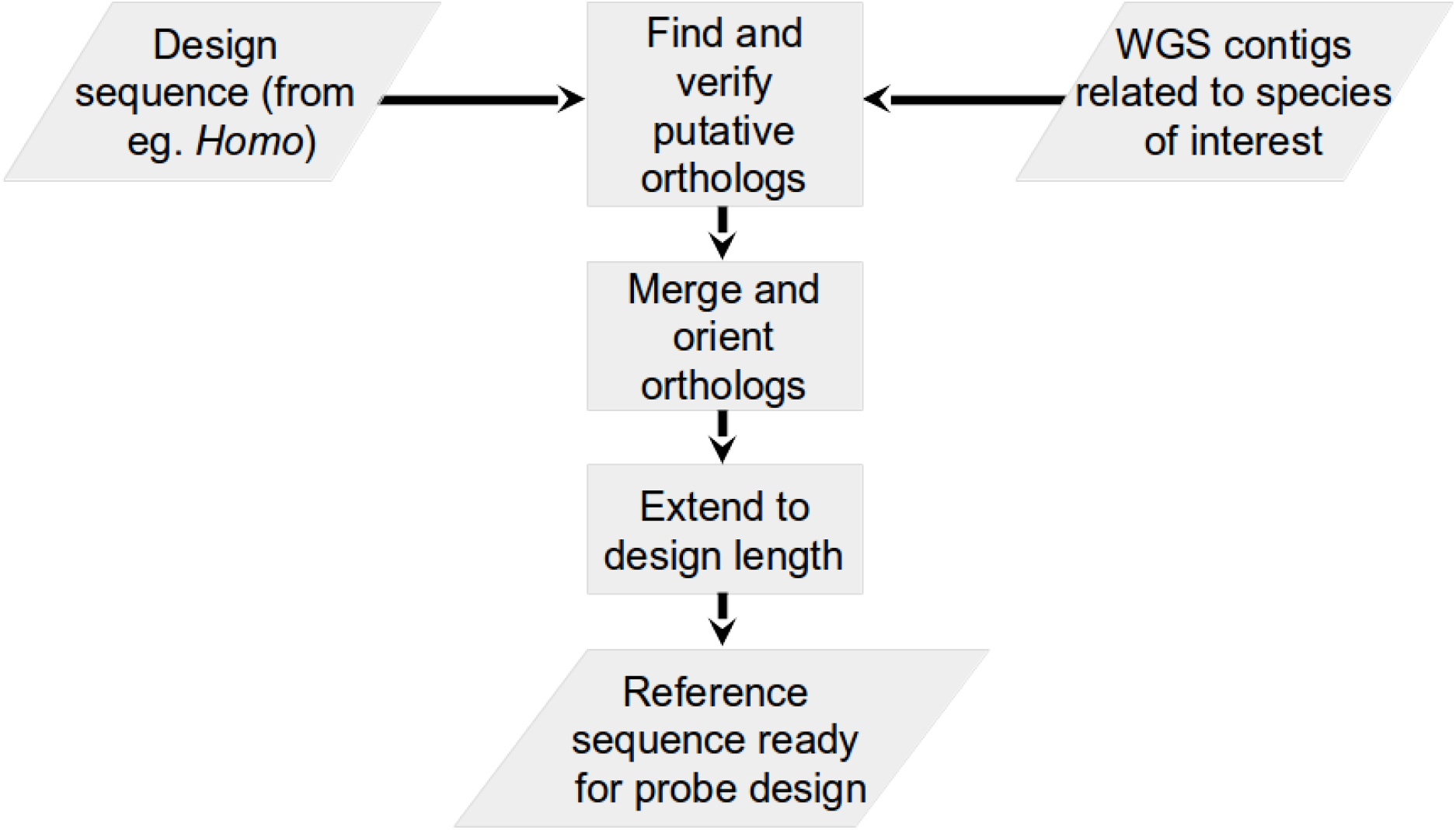
Steps of orthoCapture.

orthoCapture is intended to run on multiple WGS assemblies to maximize the diversity of the probe set. The following steps are repeated each time orthoCapture runs:

1. orthoCapture uses tblastx from BLAST (v. 2.29, Altschul 1997) on default settings to search the WGS assembly for regions similar to the exon design sequences. BLAST v. 2.29 is used in this step to retrieve nucleotide sequences because subsequent versions of BLAST output alignments as amino acid sequences rather than the necessary nucleotide sequences.
2. For each sequence retrieved from the WGS assembly, it then uses blastn from BLAST (v. 2.7.1, Camacho et al. 2009) on default settings to reciprocally match the returned sequences to the full NCBI nucleotide database, which is a collection of sequences from GenBank, RefSeq, the Third-Party Annotation (TPA) database, and the Protein Data Bank (PDB). This step verifies that the returned sequences are similar to sequences previously identified as the gene of interest.
4. After retrieving orthologous sequences from the WGS assembly, orthoCapture attempts to merge partial matches and collapse duplicates. Partial sequences are merged when they overlap perfectly by at least 30 nucleotides, and perfect matches are collapsed as duplicate records. 30 nucleotides was chosen as the default minimum overlap to minimize the number of incorrectly merged sequences (Magoc and Salzberg 2011). While paired-end read merging programs typically use a minimum overlap of 10 nucleotides (Magoc and Salzberg 2011; Liu et al. 2012; Zhang et al. 2014), this overlap length is still prone to incorrectly merging reads (Magoc and Salzberg 2011) and the longer length of WGS contigs allows a larger minimum to reduce incorrect merges that would result in poor probe design.
5. To facilitate probe design, all target sequences are then oriented as the forward or reverse complement to match the orientation of the original design sequences using seqOrient.pl (Takebayashi 2013).
6. Lastly, all target sequences are checked for minimum length. Any sequences ≤ 120 nucleotides long are extended equally on both 5’ and 3’ ends to reach this minimum probe length.

### Evaluation

orthoCapture’s reliability at discovering orthologues from WGS assemblies was evaluated by running it on WGS assemblies of species with annotated genomes. A set of 1035 random exons was generated using molbiotools Random Gene Set Generator (http://www.molbiotools.com/randomgenesetgenerator.html) and their sequences were retrieved from *Homo sapiens* based on genome annotations using BioMart from ensembl (Zerbino et al. 2018). Sequences for these genes were also retrieved from five species with annotated genomes, *Pan paniscus, P. troglodytes, Mus musculus, Rattus norvegicus*, and *Canis familiaris*. orthoCapture was run on WGS assemblies of each of these five species using *H. sapiens* design sequences of the 1035 exons. The orthologs retrieved by orthoCapture were then compared to the exons identified by annotation in the corresponding genomes (figure 3).

**Figure 3.**
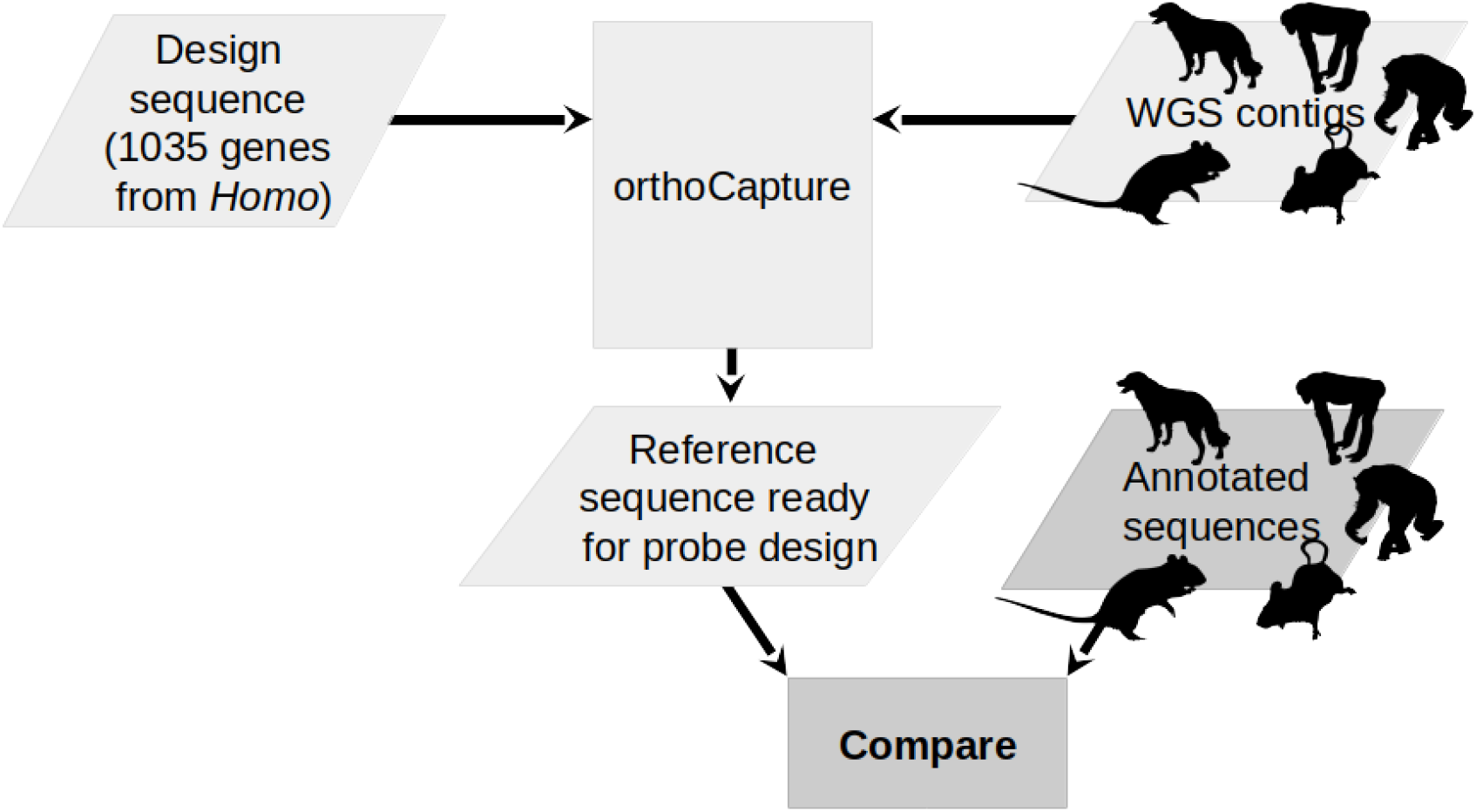
Evaluation steps for orthoCapture. Silhouettes from phylopic.org, *Canis familiaris* by Tracy A. Heath; *Mus musculus* by David Liao (CC BY-SA 3.0); *Pan troglotytes* by T. Michael Keesey (vectorization) and Tony Hisgett (photography) (CC BY-SA 3.0); *P. paniscus* by T. Michael Keesey; *Rattus norvegicus* by Rebecca Groom (CC BY-SA 3.0).

For each species, the target sequences discovered with orthoCapture were then compared to the sequences from the corresponding annotated exons, as well as from the NCBI nucleotide database. If orthoCapture discovered the correct sequence for an exon from a particular WGS assembly, the orthoCapture target sequence should match with 100% sequence identity for at least the length of the exon from the corresponding annotated genome and/or the gene as recorded for that species in the NCBI genome database. An orthoCapture-generated target sequence that is not a perfect match to, and/or shorter than the exon sequence from the annotated genome would indicate that orthoCapture returned a partial or erroneous sequence.

Target and annotation exon sequences were compared by local alignment and calculation of percent sequence identity and length using exonerate (Slater and Birney 2005). Those exon sequences retrieved by orthoCapture that were shorter than the respective annotated sequences were examined further by comparison with the design sequence. When orthoCapture retrieved target sequences not present in the genome annotation, these sequences were BLASTed against the NCBI nucleotide database (NCBI Resource Coordinators et al. 2018), as well as compared with the design sequence, and considered correct if matching sequences were identified in the database as the gene of interest.

## Evaluation Results

orthoCapture retrieved sequences for 99-100% as many exons from WGS contigs as were identified by standard annotation protocols (figure 4). The length of the sequence retrieved by orthoCapture varied by species, between 85 ± 27% (*Mus musculus*, maximum 100%) and 94 ± 17% (*Pan paniscus*, maximum 100%) (figure 5). Some exons were consistently absent across species in both orthoCapture and annotated results (figure 6), suggesting that some genes were not present in the assembly.

**Figure 4.**
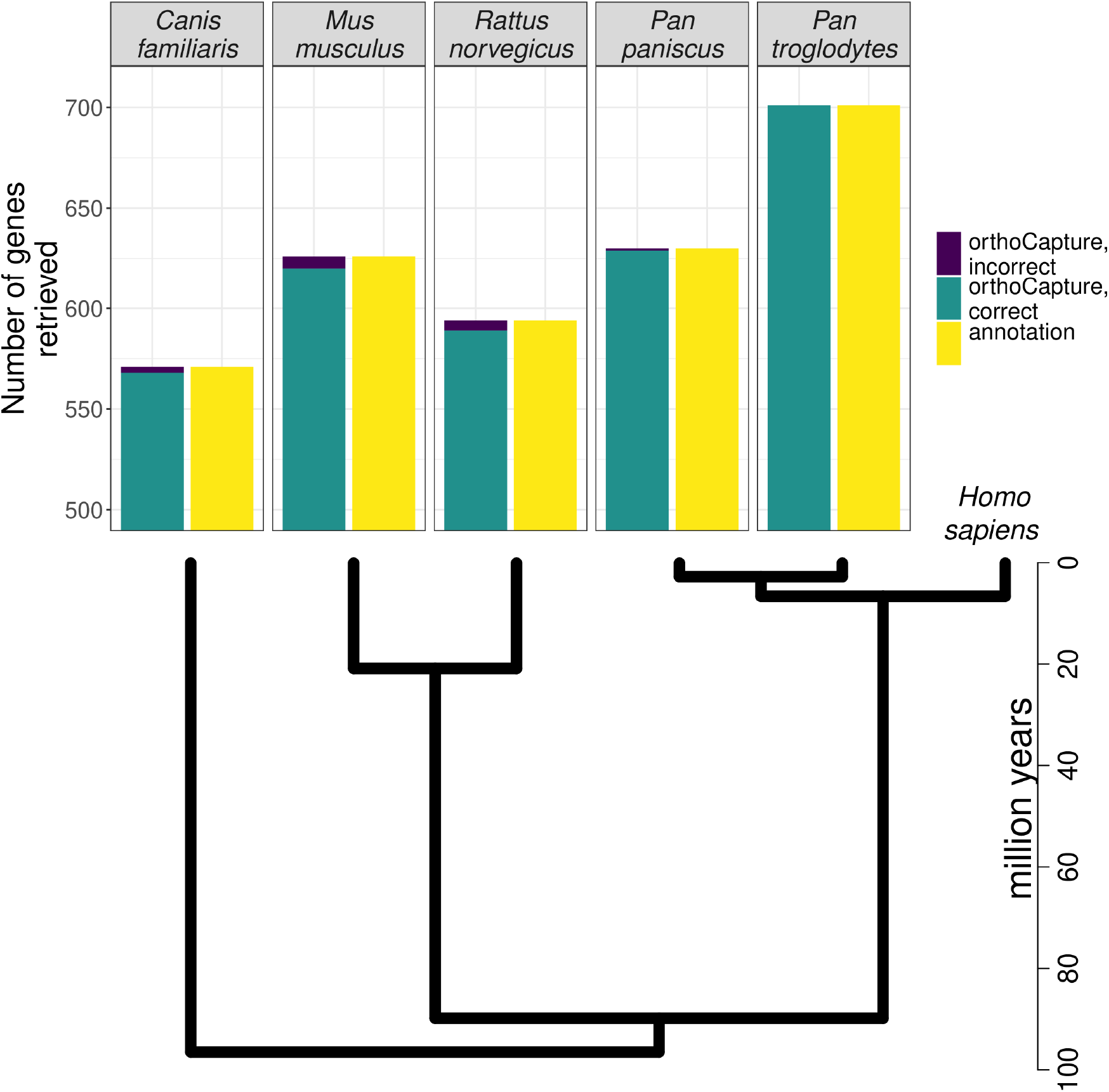
Number of exons retrieved by annotation vs orthoCapture for each genome, out of 1035 genes with annotated sequences in *Homo sapiens*. Grey bars are exons correctly retrieved by orthoCapture (true positives), stacked black bars are exons incorrectly retrieved by orthoCapture (false positives), and hatched bars are exons identified by annotation. (All exons identified by annotation are assumed to be correct.)

**Figure 5.**
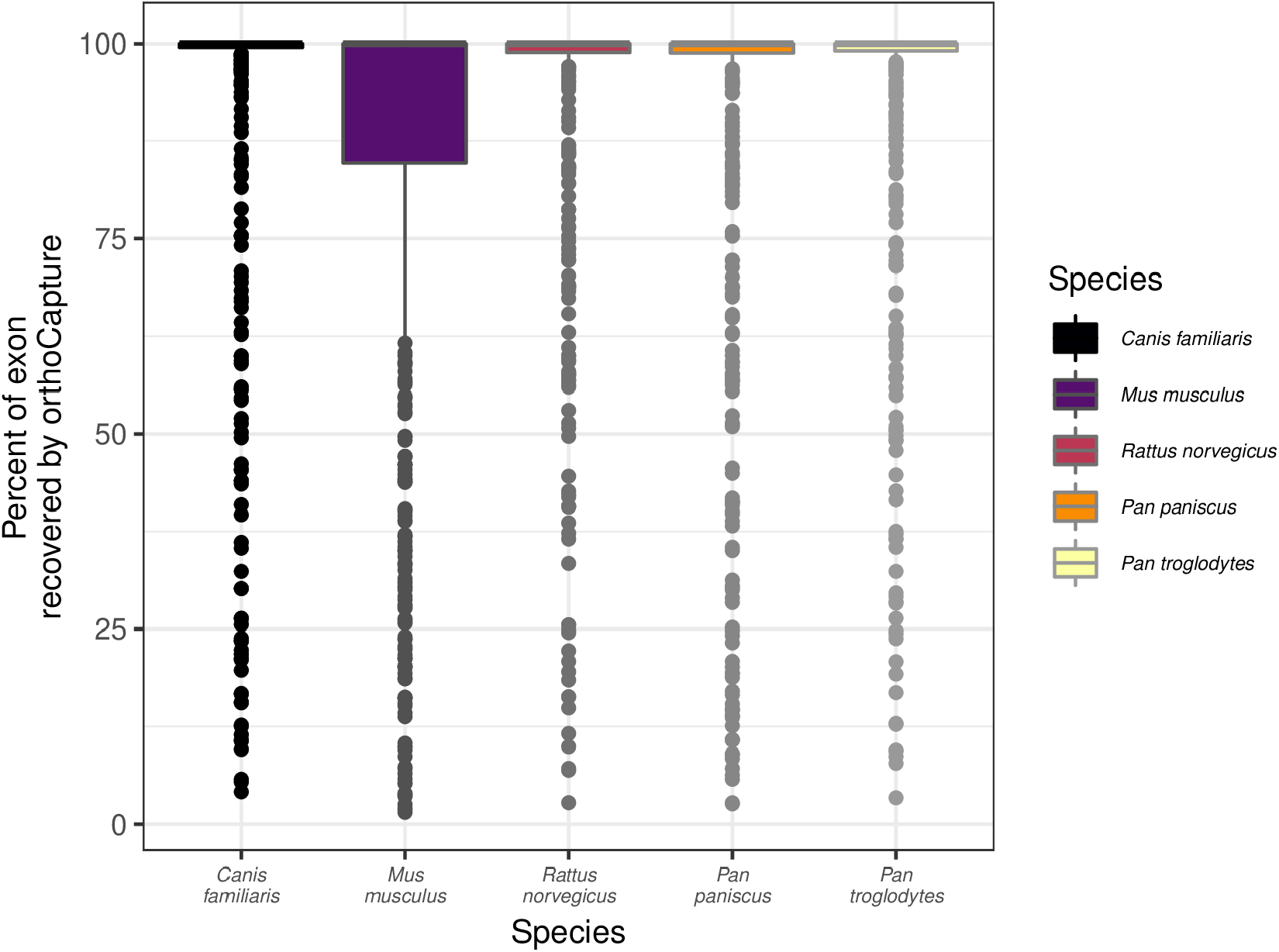
Percent of exon recovered by orthoCapture compared to annotated version. Percent recovery is shown only for those exons which were both recovered correctly by orthoCapture and annotated. Annotations were always considered to be correct and complete.

**Figure 6.**
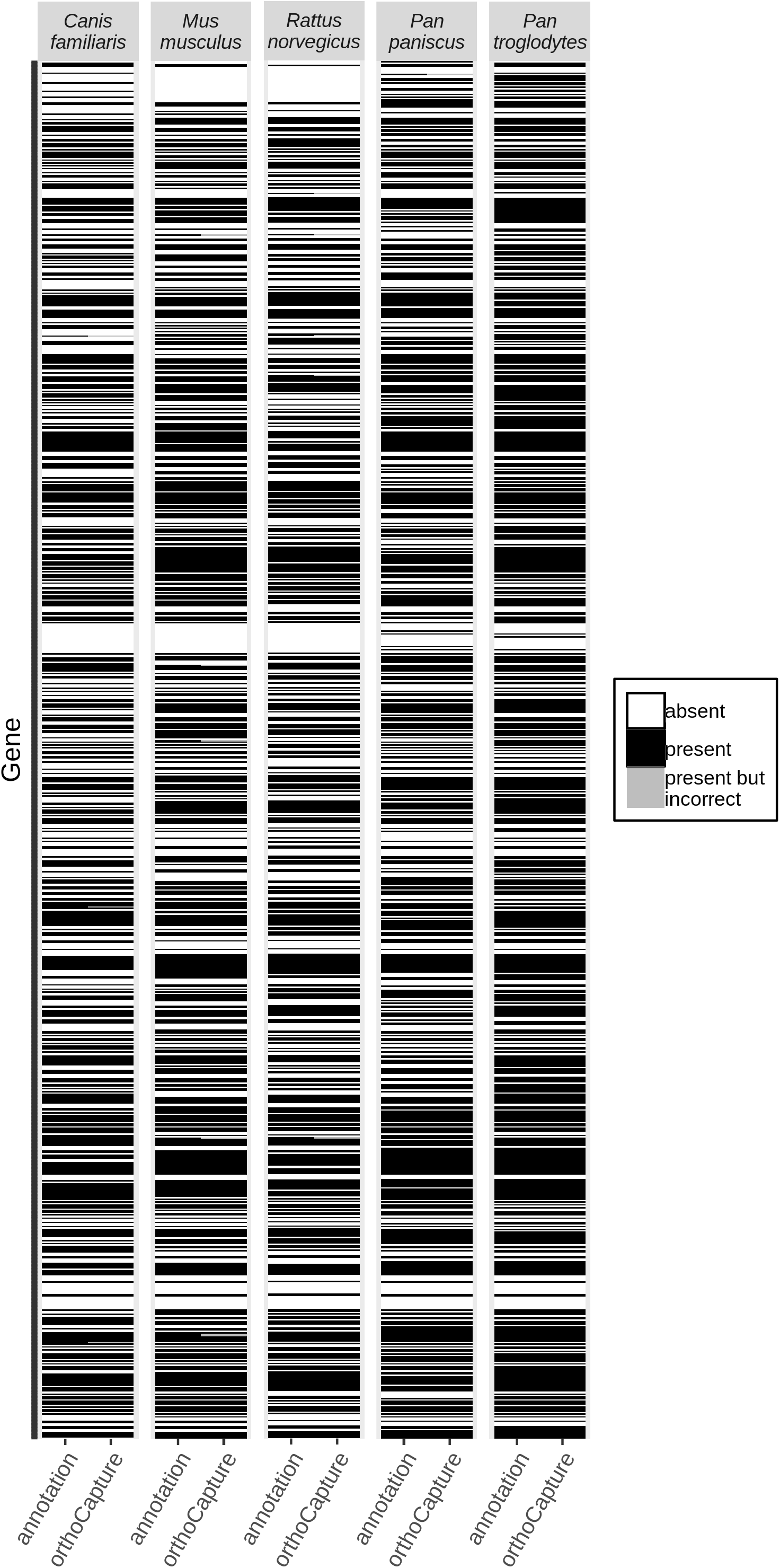
Heat maps of sequences retrieved by orthoCapture vs. from the annotation for *Homo sapiens-based* comparisons. Exons recovered by orthoCapture were considered to be correct if they had > 80% identity match with the annotated version.

The false positive rate for orthoCapture (retrieved an erroneous sequence) was less than 1% for all species, and it retrieved at least a partial sequence for all annotated exons.

## Discussion

orthoCapture reliably and accurately retrieves orthologous target sequences from whole-genome shotgun assemblies. It is capable of retrieving nearly the full length of most exons for which there are annotations. However because it is not always able to retrieve the entire exon, it is unsuitable for use as an alternative when a well-annotated genome is available. This tool makes gene capture practical for species lacking a well-annotated reference genome or one from a closely-related species.

Target capture has been a useful tool for phylogenetic analyses because it is possible to discover and design probes from (ultra-)conserved elements across diverse taxa (Bi et al. 2012; Faircloth et al. 2012; Lemmon, Emme, and Lemmon 2012; Faircloth 2017). Conserved elements are inadequate when sequence divergence is targeted, so the application of target capture has so far been limited in molecular ecology studies to species with well-annotated reference genomes or transcriptomes (or close relatives with such).

Other methods to prepare sequences for target capture probe design include HybSeq (Weitemier et al. 2014), MarkerMiner (Chamala et al. 2015), methods described by Yang and Smith (2014) and Kadlec et al. (2017), and a number of others designed for individual studies (including Ilves and López-Fernández 2014; Mandel et al. 2014; Hugall et al. 2016; Teasdale et al. 2016; Schmickl et al. 2016; Schott et al. 2017; Johnson et al. 2018). These methods routinely capture an average of 93% of target loci (Vatanparast et al. 2018). However these methods all require transcriptomes (or expressed mRNA for full exome capture, e.g., Eec-seq (Puritz and Lotterhos 2017)) from the species of interest or a close relative, which restricts their application.

orthoCapture extends target capture to species without well-annotated reference genomes or transcriptomes, thus expanding its application to taxa with few existing genomic resources that may otherwise be understudied. It does so by taking advantage of broad orthology between species of interest and species with annotated reference genomes, as well as the increased likelihood of successful capture with increased probe diversity (Bi et al. 2012; Schott et al. 2017) (figure 1).

This increased diversity presents a functional drawback to the application of orthoCapture: higher probe diversity means that more probes must be synthesized per locus, which in turn exacerbates the trade-off between number of loci covered and cost. orthoCapture would be complemented by methods such as BaitFisher (Mayer et al. 2016) or the k-medoid clustering method developed by Johnson et al. (2018), which minimize the number of probes required to capture target loci in multiple species.

### System Requirements

orthoCapture runs with a command-line interface and requires Python version 3.5 or later (see manual for required modules), as well as Perl 5.24, Bioperl, and bioperl-run. It also requires BLAST version 2.2.29 and BLAST version 2.7.1, which are linked with orthoCapture on github. A set of 1,000 exons, each less than 1,500 bp long, takes approximately 3 days running on 10 nodes with 500 gb RAM. It is recommended to break up large sets of exons into smaller batches (10-100) and run as an array if using a job scheduler (e.g., PBS or SLURM).

### Resources

Source code and a detailed manual are available at https://github.com/lauterbur/orthocapture.

## Acknowledgements

Thanks go to Liliana M. Dávalos and Krishna R. Veeramah for constructive discussions and comments. Development and testing was performed on the Stony Brook University Seawulf HPC cluster and Indiana University HPC Mason and Carbonate clusters. This research was supported, in part, by an American Association of University Women Dissertation Fellowship to M. Elise Lauterbur, and NSF-DEB1456455 to Liliana M. Dávalos.

## Data Accessibility

The data that support the evaluation are available at https://github.com/lauterbur/orthocapture. These data were derived from the following resources available in the public domain: BioMart in ensemble at https://www.ensembl.org/biomart/martview and the molbiotools Random Gene Set Generator at http://www.molbiotools.com/randomgenesetgenerator.html

## Author Contributions

M. Elise Lauterbur designed, coded, and evaluated the program, and wrote the paper.

